# Optimal fertigation for high yield and fruit quality of greenhouse strawberry

**DOI:** 10.1101/810606

**Authors:** Wu Yong, Li Li, Li Minzan, Zhang Man, Sun Hong, Nikolaos Sygrimis

**Affiliations:** Key Laboratory of Modern Precision Agriculture System Integration Research, Ministry of Education, China Agricultural University, Beijing, China; Key Laboratory of Agricultural Information Acquisition Technology, Ministry of Agriculture and Rural Areas, China Agricultural University, Beijing, China; Department of Agricultural Engineering, Agricultural University of Athens, Athens, Greece

## Abstract

Nitrogen (N), phosphorus (P), potassium (K), and water are four crucial factors that have significant effects on strawberry yield and fruit quality. A quadratic regression orthogonal rotation combination experiment that involved 36 treatments with five levels of the four variables (N, P, and K fertilizers and water) was executed to optimize the fertilization and water combination for high yield and quality. SSC/TA ratio (the ratio of soluble solid content to titratable acid) was selected as the index of quality. Results showed that the N fertilizer was the most important factor, followed by water and P fertilizer, and the N fertilizer had a significant effect on yield and SSC/TA ratio. By contrast, the K fertilizer had a significant effect only on yield. N×K fertilizer interaction had a significant effect on yield, whereas the other interactions among the four factors had no significant effects on yield and SSC/TA ratio. The effects of the four factors on the yield and SSC/TA ratio were ranked as N fertilizer > water > K fertilizer > P fertilizer and N fertilizer > P fertilizer > water > K fertilizer, respectively. The yield and SSC/TA ratio increased and then decreased when NPK fertilizer and water increased. The optimal fertilizer and water combination was 22.28–24.61 g/plant Ca (NO_3_)_2_⋅4H_2_O, 1.75–2.03 g/plant NaH_2_PO_4_, 12.41–13.91 g/plant K_2_SO_4_, and 12.00– 13.05 L/plant water for yields of more than 110 g/plant and optimal SSC/TA ratio of 8.5–14.

## Introduction

Mineral fertilizers and water have a significant effect on crop yield [1–3]. However, the excessive application of fertilizers may lead to soil and water pollution and become a serious threat to food safety [4,5]. Meanwhile, water scarcity is now a major challenge in China [6]. Therefore, a good management of fertilization and water is increasingly required for agriculture in China.

Strawberry is one of the most profitable fruit cultivars in China, which ranks first in total strawberry production worldwide with a production of 1,801,865 tons in 2016, followed by United States and Mexico among a total of 79 countries [7]. Thus, large amounts of fertilizers and water are needed for strawberry production in China. Consumers prefer strawberries with a sweet taste [8,9]. The strawberry flavor is strongly correlated with the balance between the soluble solid content (SSC) and titratable acid (TA) in ripe fruits [10,11], which are common quality indexes to assess sweetness and sourness [12]. The ratio of SSC/TA is another effective parameter to determine fruit flavor [13,14]. The higher the SSC/TA ratio, the sweeter the fruit [15]. Therefore, increasing strawberry production and enhancing fruit quality with high SSC/TA ratio and without environment pollution are important goals in the field.

Nitrogen (N), phosphorus (P), and potassium (K) are primary mineral fertilizers. N is the most limiting nutrient to crop production because of its important role in cell division [16,17], and N deficiency can decrease crop yield and quality [18,19]. P nutrient is essential for photosynthesis [20], and it is required after emergence [21]. K is the second most abundant element in plant tissues after N [22,23], and it helps enhance water uptake and grain quality [24]. In addition to mineral fertilizers, water greatly contributes to the strawberry fruit content and leaf development [25], and water shortages can lead to large losses of strawberries yield [26].

Although studies have shown that all mineral fertilizers (N, P, and K) and water have effects on the yield and quality of strawberries, most of them only focused on either the effect of water [26–28] or the effect of fertilization [29–33]. The combined application of N, P, and K (NPK) fertilizers and water for high yield and good fruit quality is rarely reported [27,31,34]. Thus, the interaction effect among N, P, and K fertilizers and water on the strawberry yield and fruit quality should be investigated. This study aimed to evaluate the influence of different levels of NPK fertilizers and water on the growth and fruit quality of strawberry and determine the optimal fertilization and water combination for high yield and good quality.

## Materials and methods

### Experimental site and cultivar

The experiment was conducted for 8 months from November 2016 to June 2017 in an east–west oriented solar greenhouse located in the Zhuozhou Experiment Center, China Agricultural University, China. The Chinese solar greenhouse, as a horticultural facility, is a kind of mono-slope greenhouse that provides effective energy use and is widely used in China, especially in the northern latitudes [35,36]. The structure of this solar greenhouse has a typical width, length, backwall height, and roof height of 8, 50, 2.4, and 3.5 m, respectively (Fig 1).

**Fig 1.**
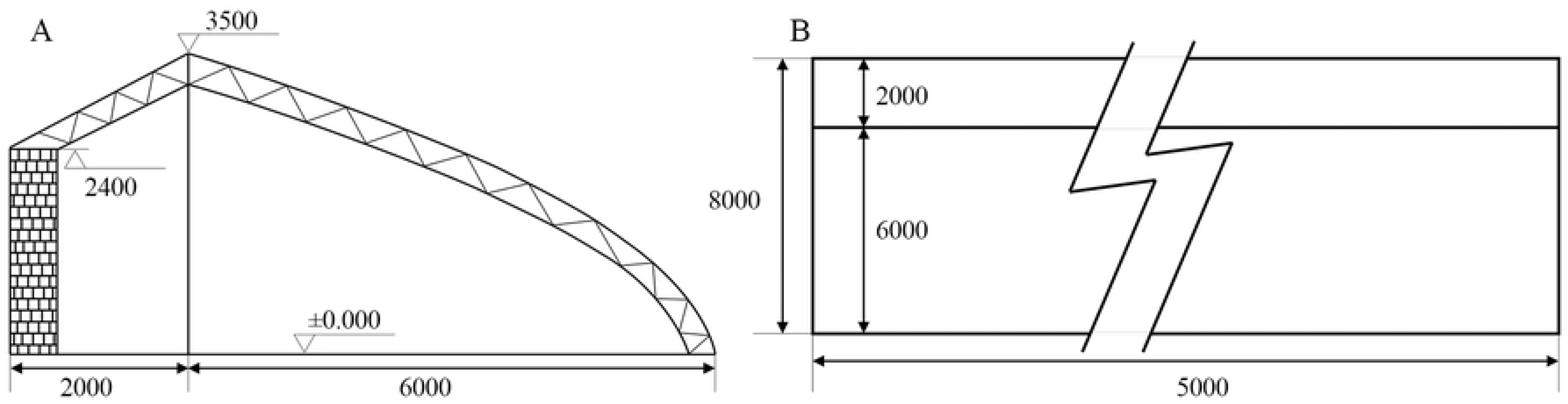
Solar greenhouse block diagram. (A) Section view. (B) Top view.

The strawberry cultivar used was ‘Hongyan’, which has been studied extensively in China [37–40]. It was cultivated on substrate instead of soil in the solar greenhouse with natural light and temperature of 10 °C–26 °C. The substrate, with initial chemical characteristics shown in Table 1, was a mixture with a ratio of 10:2:1 rate of peat, vermiculite, and perlite, respectively. The substrate bags, with dimensions of 100 cm×40 cm×20 cm, were obtained from Beijing Greenovo Agriculture Science and Technology Co., Ltd., and every three strawberry plants were grown on each substrate bag (Fig 2). The strawberries were transplanted on November 3, 2016 and hand-harvested at the mature red stage. Thereafter, the fruits were transported to the laboratory within 2 h by using ice bags for cooling.

**Table 1.** Substrate chemical analysis (%)

| Properties | Fertility |
| --- | --- |
| Total Nitrogen | 0.9705 |
| Total Phosphorus | 0.466 |
| Potassium | 1.07 |

**Fig 2.**
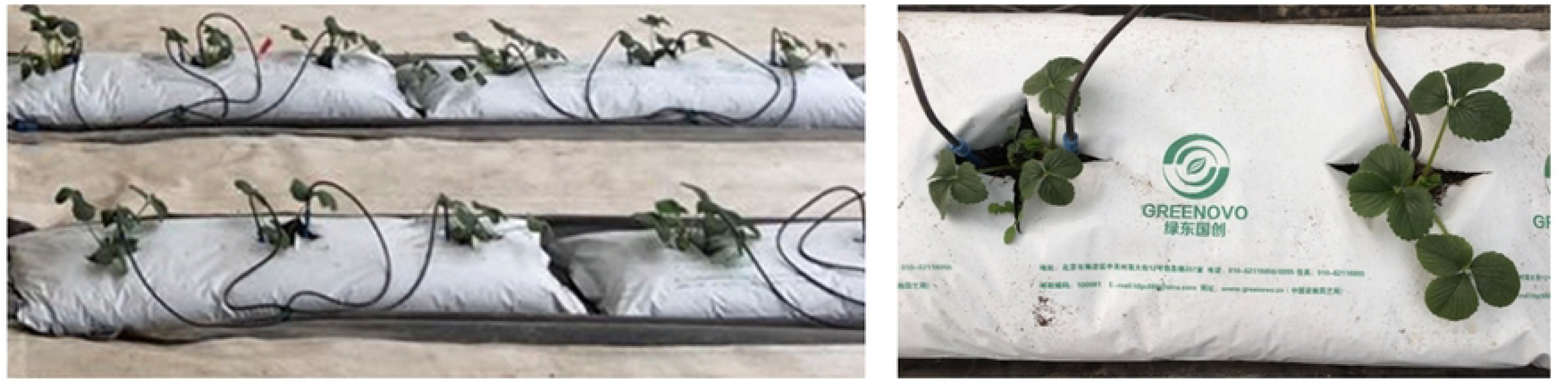
Strawberry plants grown on substrate bags.

### Experimental method

To reveal the relationship between the NPK+water combination and the fruit yield and achieve the optimal combination, a quadratic regression orthogonal rotation combination experiment was designed involving four factors at five levels in 36 treatments; this technique is currently the most effective method for multi-factor interaction effect analysis [41]. The four factors were N, P, and K fertilizers and water, which were represented by *x*_*1*_, *x*_*2*_, *x*_*3*_, and *x*_*4*_, respectively. Calcium nitrate (Ca (NO_3_)_2_⋅4H_2_O), sodium dihydrogen phosphate (NaH_2_PO_4_), and potassium sulfate (K_2_SO_4_), which were obtained from Shanghai Wintong Chemicals Co., Ltd. with a purity of more than 99%, were used as the sources of N, P, and K, respectively. Tap water was used as the source of water. The arrangement of these factors and the levels of variables chosen are shown in Table 2. All treatments were arranged in a completely randomized block with three replications, and each treatment consisted of six plants. A total of 648 plants grown in the experimental solar greenhouse were studied. All plants were fed with a mixture solution of the NPK fertilizers and water weekly following each treatment arrangement 15 days after the transplant, and additional macronutrients and micronutrients were also applied weekly with the same dosage for each treatment (Table 3). The treatment details are presented in Table 4.

**Table 2.** Design level of four variables in the quadratic regression orthogonal experiment

| Variable X | Changing interval $\Delta_j$ | Design level of variables ($m_0 = 12, \gamma = 2$) | | | | |
| --- | --- | --- | --- | --- | --- | --- |
| | | $-\gamma$ | $-1$ | $0$ | $+1$ | $+\gamma$ |
| $x_1$ (g/plant) | 13.41 | 12.57 | 16.77 | 20.96 | 25.15 | 29.34 |
| $x_2$ (g/plant) | 1.21 | 1.14 | 1.52 | 1.89 | 2.27 | 2.65 |
| $x_3$ (g/plant) | 4.78 | 4.48 | 5.98 | 7.48 | 8.97 | 10.47 |
| $x_4$ (L/plant) | 7.68 | 7.20 | 9.60 | 12.00 | 14.40 | 16.80 |

**Table 3.** Additional macronutrients and micronutrients applied to each treatment with the same dosage

| Fertilizers | Dosage (g/plant) |
| --- | --- |
| $\text{MgSO}_4$ | 4.43 |
| EDTA-2NaFe | 3.5461 |
| $\text{H}_3\text{BO}_3$ | 0.3381 |
| $\text{MnSO}_4 \cdot 4\text{H}_2\text{O}$ | 0.2518 |
| $\text{ZnSO}_4 \cdot 7\text{H}_2\text{O}$ | 0.0260 |
| $\text{CuSO}_4 \cdot 5\text{H}_2\text{O}$ | 0.0095 |
| $(\text{NH}_4)_6\text{Mo}_7\text{O}_{24} \cdot 4\text{H}_2\text{O}$ | 0.0024 |

**Table 4.**
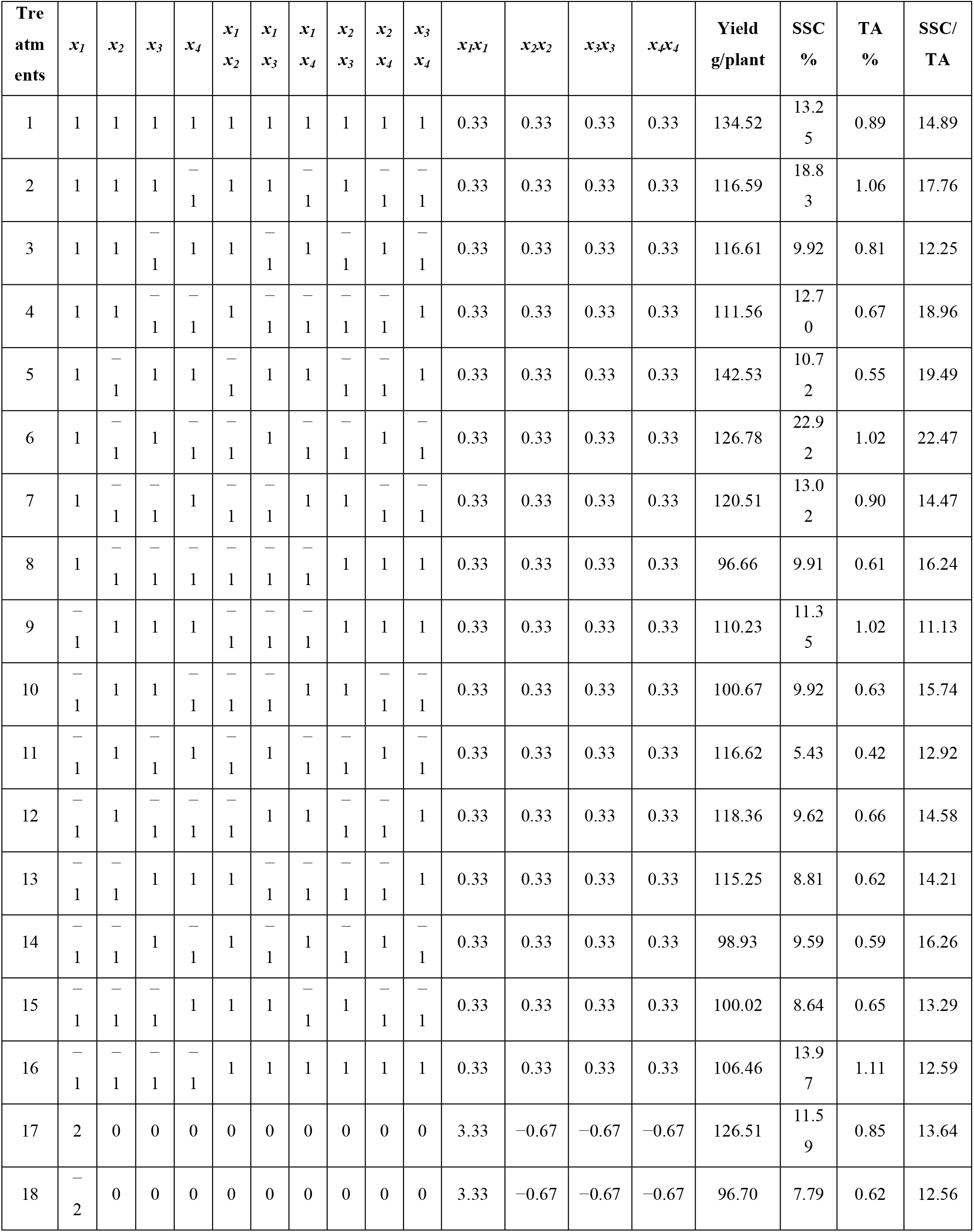
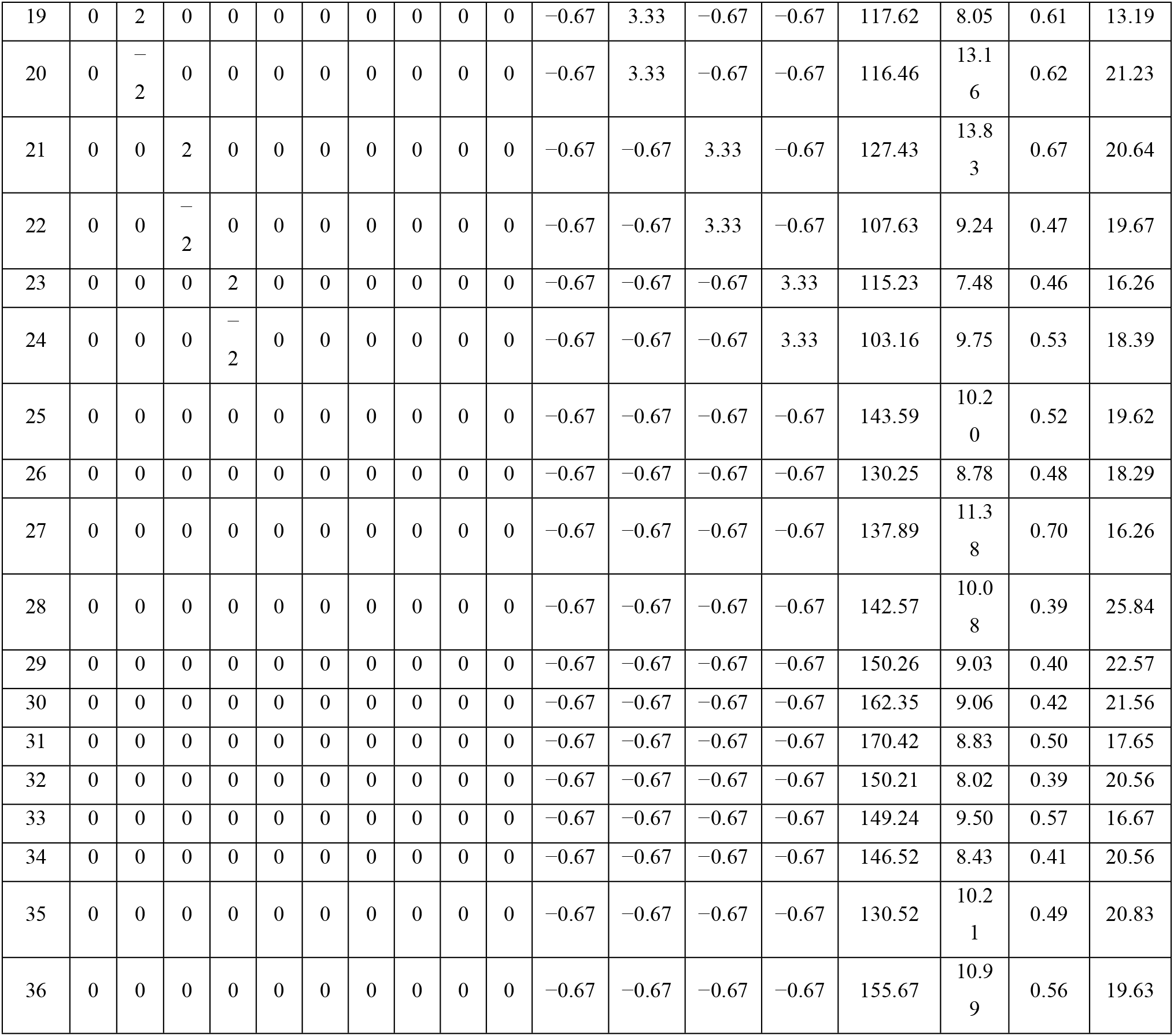
Arrangement of variables in the quadratic regression orthogonal experiment and results of the experiment

### Measurement of yield and fruit quality traits

An analytical balance (0.01 g accuracy) was used to measure the fruit weight after the fruits were harvested. Parameters of the fruit quality, namely, soluble solid content (SSC) and titratable acidity (TA), were measured after the fruits were transported to the laboratory. SSC (%) was determined by a digital hand-held pocket refractometer (PAL-1, Atago, Japan), whereas TA (%) was measured by neutralization to pH 7.0 with 0.1 N NaOH. Data were presented as percentages of malic acid.

## Results

### Yield respond to N, P, and K fertilizers and water

After a significance test on regression coefficients and regression formulas, the equation that governs the effect of N (*x*_*1*_), P (*x*_*2*_), K (*x*_*3*_), and water (*x*_*4*_) on yield (*y_1_*) is formulated as follows:

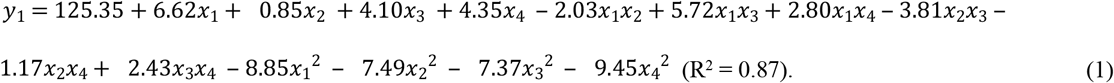

ANOVA results are shown in Table 5. The regression was significant at the 0.01 probability (F = 10.29 > F_0.01_ (14, 21) = 3.07), indicating that the regression model was a good fit for the experimental data. The F-value for the lack-of-fit test was 0.13. This value was less than the significant value at the 0.05 probability (F_0.05_ (10, 11) = 2.85), which was insignificant; the regression model was relatively suitable. Therefore, this regression model could be used to evaluate the effects of N, P, and K fertilizers on the ‘Hongyan’ strawberry yield.

**Table 5.** ANOVA table of effects of N (x1), P (x2), K (x3), and water (x4) on yield (y1).

| Source of variance | df | SS | MS | F-value | Significance |
| --- | --- | --- | --- | --- | --- |
| $x_1$ | 1.00 | 1051.26 | 1051.26 | 12.74 | ** |
| $x_2$ | 1.00 | 17.24 | 17.24 | 0.21 | ns |
| $x_3$ | 1.00 | 402.62 | 402.62 | 4.88 | * |
| $x_4$ | 1.00 | 454.31 | 454.31 | 5.51 | * |
| $x_1x_2$ | 1.00 | 65.69 | 65.69 | 0.80 | ns |
| $x_1x_3$ | 1.00 | 522.81 | 522.81 | 6.34 | * |
| $x_1x_4$ | 1.00 | 125.89 | 125.89 | 1.53 | ns |
| $x_2x_3$ | 1.00 | 232.41 | 232.41 | 2.82 | ns |
| $x_2x_4$ | 1.00 | 21.81 | 21.81 | 0.26 | ns |
| $x_3x_4$ | 1.00 | 94.28 | 94.28 | 1.14 | ns |
| $x_1^2$ | 1.00 | 2506.56 | 2506.56 | 30.38 | ** |
| $x_2^2$ | 1.00 | 1796.00 | 1796.00 | 21.77 | ** |
| $x_3^2$ | 1.00 | 1737.75 | 1737.75 | 21.06 | ** |
| $x_4^2$ | 1.00 | 2859.44 | 2859.44 | 34.66 | ** |
| Regression | 14.00 | 11888.07 | 849.15 | 10.29 | ** |
| Residual | 21.00 | 1732.65 | 82.51 |  |  |
| Lack of fit | 10.00 | 183.29 | 18.33 | 0.13 | ns |
| Error | 11.00 | 1549.35 | 140.85 |  |  |
| Total | 35.00 | 13620.72 |  |  |  |

$F_{0.05}(1, 21) = 4.32$ , $F_{0.01}(1, 21) = 8.02$ , $F_{0.05}(14, 21) = 2.20$ , $F_{0.01}(14, 21) = 3.07$ , $F_{0.05}(14, 21) = 2.20$ , $F_{0.05}(10, 11) = 2.85$ ,
$F_{0.01}(10, 11) = 4.54$ .
\* and \*\* are significant at the 0.05 and 0.01 probability levels, respectively.
ns: non-significant differences.

As shown in Table 5, N, K, and water had a significant effect on strawberry yield, but P had no significant effect. The relative magnitude of the effects of N, P, K, and water on yield was N>water>K>P in accordance with the absolute value of the standardized regression coefficient. No significant interaction occurred between N×P, N×water, P×K, P×water, and K×water. Thus, an ideal fit equation could be obtained as follows:

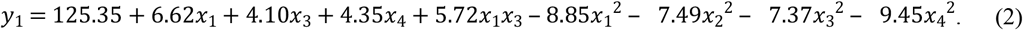

From the equation above, the partial regression equations were as follows:

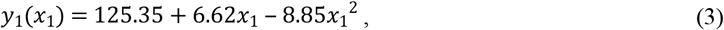

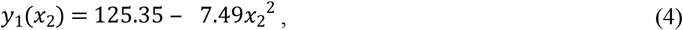

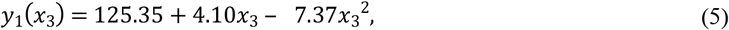

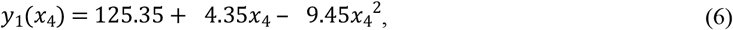

The partial regression equation results showed that yield rapidly increased with an increase in N and P fertilizers at levels below 0.37 (22.51 g/plant) and 0 (1.89 g/plant), respectively, and rapidly decreased at levels above them (Fig 3). With increasing K fertilizer, the yield rapidly increased and then slowly decreased, and it peaked at the 0.28 (7.90 g/plant) level of K fertilizer. When increasing water, the yield rapidly increased and then gradually decreased, and the maximum value was at the 0.23 (12.55 L/plant) level of water.

**Fig 3.**
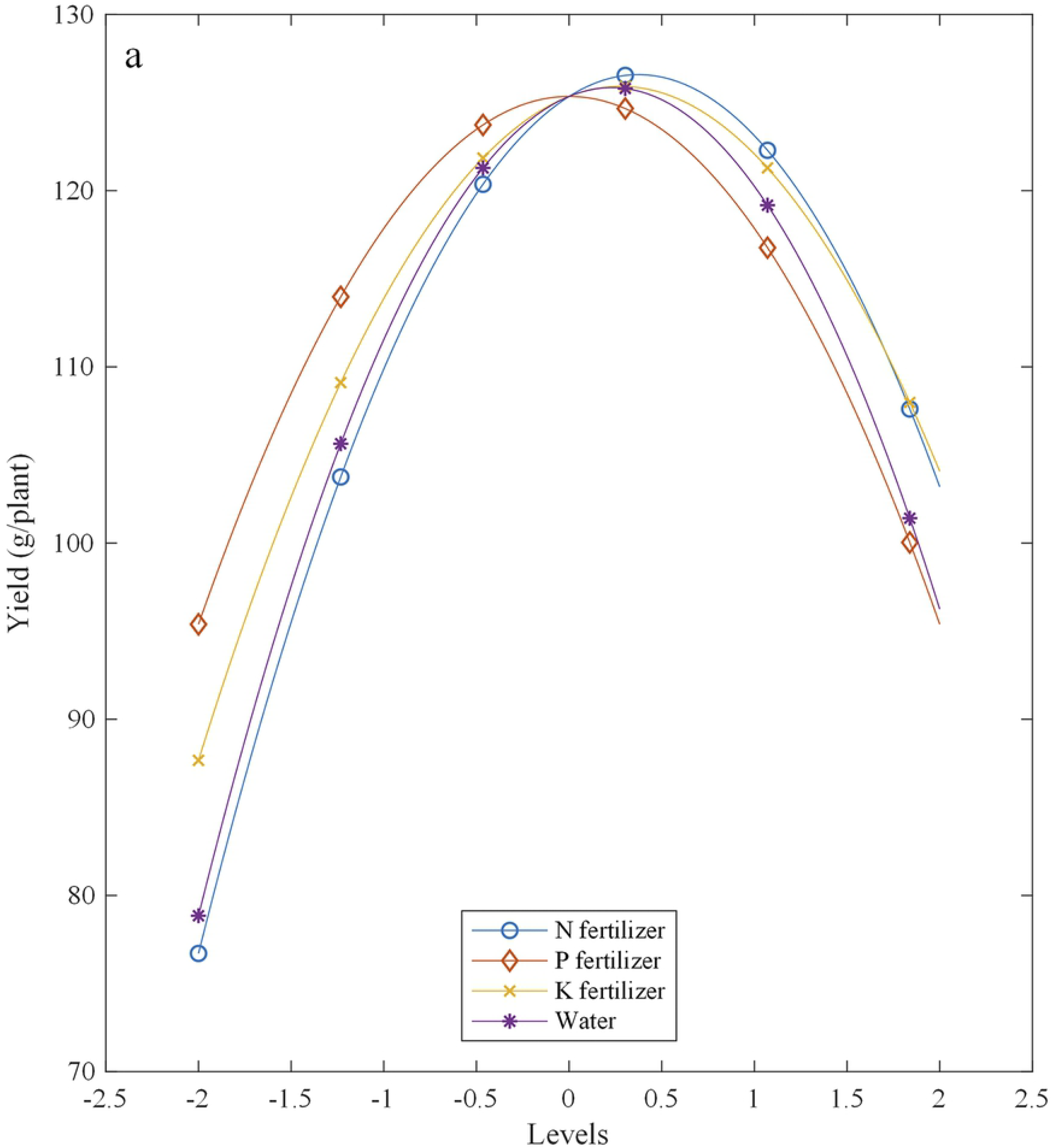
Effects of N, P, and K fertilizers and water on the yield.

The interaction effect of every two factors on yield and SSC/TA ratio are shown in Fig 4. Interaction analysis showed that yield rapidly increased and then slowly decreased as the N fertilizer increased but slowly increased and then slowly decreased as the P fertilizer increased. Furthermore, the maximum yield was 126.59 g/plant at 22.51 g/plant Ca (NO_3_)_2_⋅4H_2_O and 1.89 g/plant NaH_2_PO_4_ (Fig 4(a)). Yield rapidly increased and then slowly decreased with increasing levels of N and K fertilizers, and the maximum yield was 127.16 g/plant at 22.51 g/plant Ca (NO_3_)_2_⋅4H_2_O and 7.90 g/plant K_2_SO_4_ (Fig 4(b)). The same trends were obtained for N×water interaction, that is, the yield rapidly increased, slowly decreased, and reached the maximum yield of 127.09 g/plant at 22.51 g/plant Ca (NO_3_)_2_⋅4H_2_O and 12.55 L/plant water (Fig 4(c)). Similarly, for the P×K (Fig 4(d)), P×water (Fig 4(e)), and K×water (Fig 4(f)) interactions, the yield increased and then decreased, and the maximum yields were 125.92 g/plant at 1.89 g/plant NaH_2_PO_4_ and 7.90 g/plant K_2_SO_4_ (Fig 4(d)), 125.85 g/plant at 1.89 g/plant NaH_2_PO_4_ and 12.55 L/plant water (Fig 4(e)), and 126.42 g/plant at 7.90 g/plant K_2_SO_4_ and 12.55 L/plant water (Fig 4(f)), respectively.

**Fig 4.**
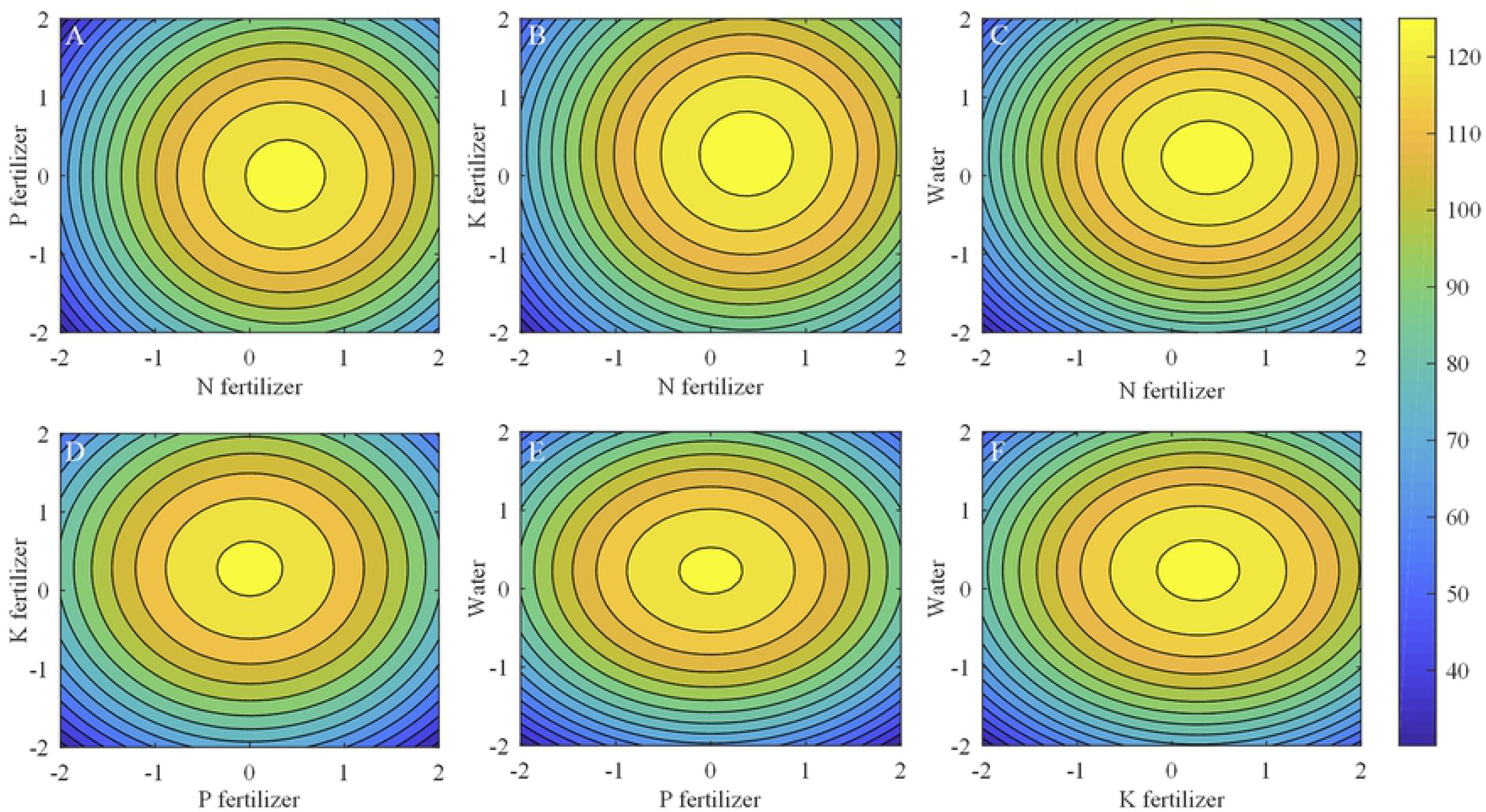
Effects of interaction among N, P, and K fertilizers and water on the yield. (A) N-P interaction effect on yield with K fertilizer and water at 0 level. (B) N-K interaction effect on yield with P fertilizer and water at 0 level. (C) N-water interaction effect on yield with P fertilizer and K fertilizer at 0 level. (D) P-K interaction effect on yield with N fertilizer and water at 0 level. (E) P-water interaction effect on yield with N fertilizer and K fertilizer at 0 level. (F) K-water interaction effect on yield with N fertilizer and P fertilizer at 0 level

Frequency analysis was conducted to obtain the optimal fertilization combination for high yield (Table 6). Among 625 kinds of fertilization combinations, 27 combinations of the four factors had a yield of more than 110 g/plant. The 99% confidence interval for N, P, and K fertilizers and water levels were 0.314–0.871, −0.357–0.357, 0.169–0.794, and 0.000–0.592, respectively. Therefore, when applying 22.28–24.61 g/plant Ca (NO_3_)_2_⋅4H_2_O, 1.75–2.03 g/plant NaH_2_PO_4_, 12.41–13.91 g/plant K_2_SO_4_, and 12.00–13.42 L/plant water, fruit yield will reach more than 110 g/plant with a probability of 99%.

**Table 6.** Frequency analysis of N, P, and K fertilizers and water for strawberry yield of more than 110 g/plant.

| Levels | N fertilizer |  | P fertilizer |  | K fertilizer |  | Water |  |
| --- | --- | --- | --- | --- | --- | --- | --- | --- |
|  | Sets Times | Frequency | Sets Times | Frequency | Sets Times | Frequency | Sets Times | Frequency |
| –2 | 0 | 0.00 | 0 | 0.00 | 0 | 0.00 | 0 | 0.00 |
| –1 | 0 | 0.00 | 7 | 0.26 | 1 | 0.04 | 2 | 0.07 |
| 0 | 12 | 0.44 | 13 | 0.48 | 13 | 0.48 | 15 | 0.56 |
| 1 | 14 | 0.52 | 7 | 0.26 | 12 | 0.44 | 10 | 0.37 |
| 2 | 1 | 0.04 | 0 | 0.00 | 1 | 0.04 | 0 | 0.00 |
| Total | 27 | 1 | 27 | 1 | 27 | 1 | 27 | 1 |
| Average | 0.593 |  | 0.000 |  | 0.481 |  | 0.296 |  |
| Standard error | 0.108 |  | 0.139 |  | 0.121 |  | 0.115 |  |
| 99% confidence interval | 0.314–0.871 |  | –0.357–0.357 |  | 0.169–0.794 |  | 0.000–0.592 |  |
| Optimal Fertilization (g/plant) | 22.28–24.61 |  | 1.75–2.03 |  | 12.41–13.91 |  | 12.00–13.42 |  |

### SSC/TA ratio responds to N, P, and K fertilizers and water

The regression equation that governs the effect on the SSC/TA ratio (*y_2_*) by N (*x*_*1*_), P (*x*_*2*_), K (*x*_*3*_), and water (*x*_*4*_) is formulated as follows:

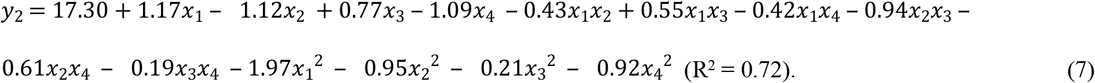

ANOVA results are shown in Table 7. The F-value for the regression model was 3.82, which was larger than F_0.01_ (14, 21) = 3.07. Thus, the regression model was a good fit for the experimental data. The F-value for the lack-of-fit test was 0.66. This value was less than the significant value at the 0.05 probability (F_0.05_ (10, 11) = 2.85), which was insignificant. Thus, the regression model was relatively suitable. This regression model could be used to evaluate the effects of N, P, and K fertilizers on the SSC/TA ratio of ‘Hongyan’ strawberry fruits.

**Table 7.** ANOVA table of effect of N, P, and K fertilizers and water on the SSC/TA ratio.

| Source of variance | *df* | *SS* | *MS* | *F-value* | *Significance* |
| --- | --- | --- | --- | --- | --- |
| $x_1$ | 1.00 | 32.60 | 32.60 | 5.46 | * |
| $x_2$ | 1.00 | 30.08 | 30.08 | 5.04 | * |
| $x_3$ | 1.00 | 14.40 | 14.40 | 2.41 | ns |
| $x_4$ | 1.00 | 28.62 | 28.62 | 4.80 | * |
| $x_1x_2$ | 1.00 | 2.92 | 2.92 | 0.49 | ns |
| $x_1x_3$ | 1.00 | 4.76 | 4.76 | 0.80 | ns |
| $x_1x_4$ | 1.00 | 2.81 | 2.81 | 0.47 | ns |
| $x_2x_3$ | 1.00 | 14.12 | 14.12 | 2.37 | ns |
| $x_2x_4$ | 1.00 | 5.94 | 5.94 | 1.00 | ns |
| $x_3x_4$ | 1.00 | 0.59 | 0.59 | 0.10 | ns |
| $x_1^2$ | 1.00 | 124.81 | 124.81 | 20.92 | ** |
| $x_2^2$ | 1.00 | 28.72 | 28.72 | 4.81 | ** |
| $x_3^2$ | 1.00 | 1.43 | 1.43 | 0.24 | ns |
| $x_4^2$ | 1.00 | 27.01 | 27.01 | 4.53 | * |
| Regression | 14.00 | 318.81 | 22.77 | 3.82 | ** |
| Residual | 21.00 | 125.31 | 5.97 |  |  |
| Lack of fit | 10.00 | 47.05 | 4.70 | 0.66 | ns |
| Error | 11.00 | 78.26 | 7.11 |  |  |
| Total | 35.00 | 456.55 |  |  |  |

$F_{0.05}(1, 21) = 4.32$ , $F_{0.01}(1, 21) = 8.02$ , $F_{0.05}(14, 21) = 2.20$ , $F_{0.01}(14, 21) = 3.07$ , $F_{0.05}(14, 21) = 2.20$ , $F_{0.05}(10, 11) = 2.85$ , $F_{0.01}(10, 11) = 4.54$ .
\* and \*\* are significant at the 0.05 and 0.01 probability levels, respectively.
ns: non-significant differences.

The N and P fertilizers and water had a significant effect on strawberry fruit’s SSC/TA ratio, but the K fertilizer had no significant effect. The relative magnitude of the effects of N, P, and K fertilizers and water on the SSC/TA ratio was N>P>water>K, which was in accordance with the absolute value of the standardized regression coefficient. No significant interaction occurred among N, P, K, and water in terms of the SSC/TA ratio (Table 7). Therefore, an ideal fit equation could be obtained as follows:

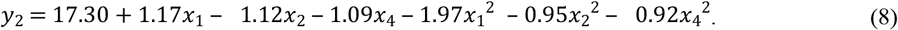

From the equation above, the partial regression equations were as follows:

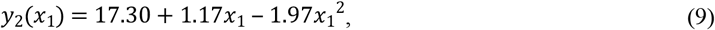

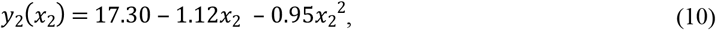

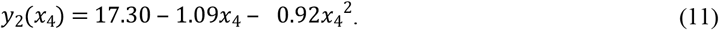

The partial regression equation results showed that the SSC/TA ratio rapidly increased with an increase in P and water at levels below −0.59 (1.67 g/plant) and −0.59 (10.58 g/plant), respectively, and slowly decreased at levels above them (Fig 5). With increasing N, the SSC/TA ratio gradually increased and then gradually decreased, with a maximum value at the 0.30 (22.22 g/plant) level of N.

**Fig 5.**
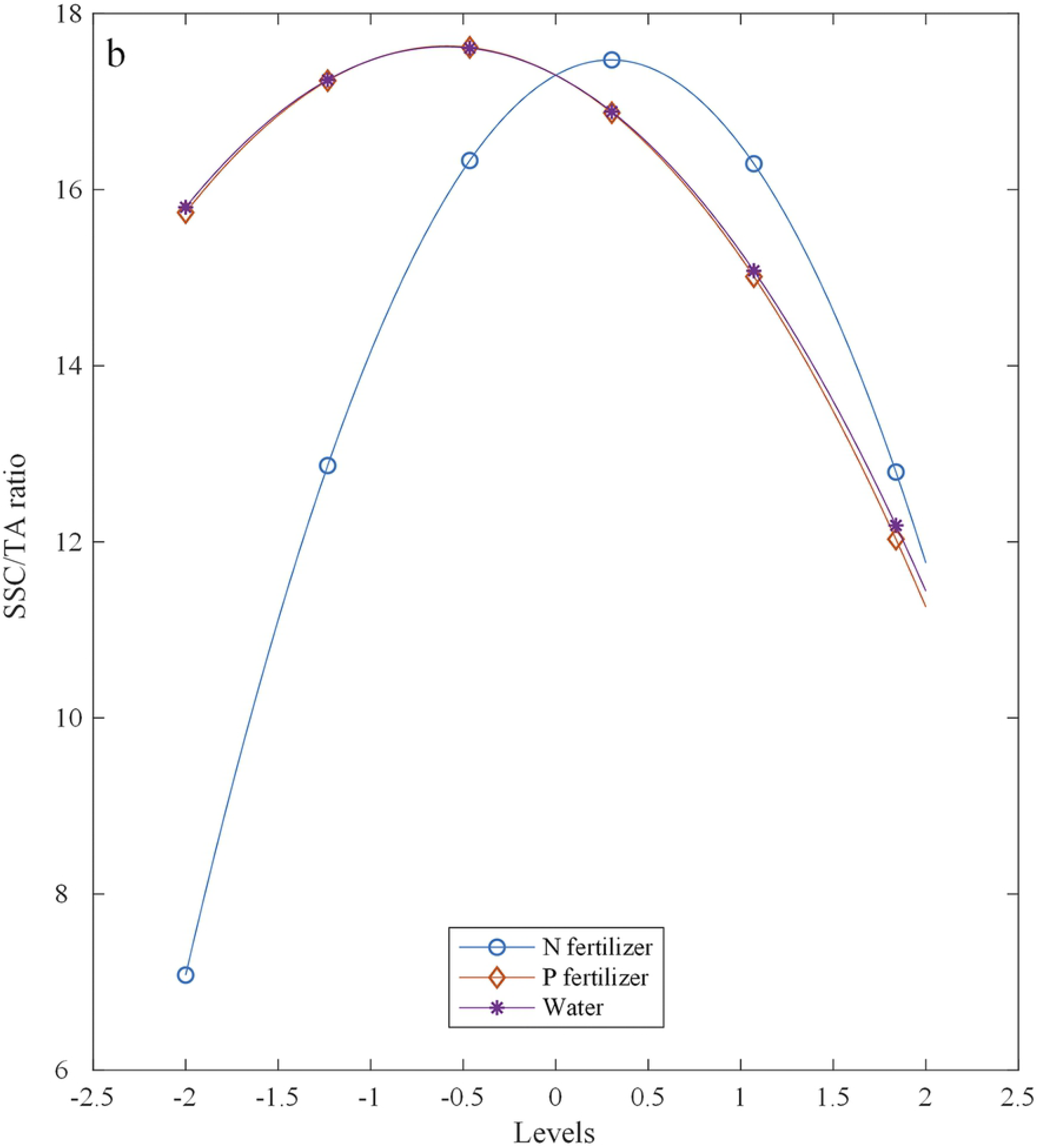
Effects of N, P, and K fertilizers and water on the SSC/TA ratio.

The SSC/TA ratio rapidly increased and then rapidly decreased with increasing levels of N, whereas it slowly increased and then rapidly decreased with increasing P; meanwhile, the maximum SSC/TA ratio was 17.80 at 22.22 g/plant Ca (NO_3_)_2_⋅4H_2_O and 1.67 g/plant NaH_2_PO_4_ (Fig 6(a)). The same trend in Fig 6(a) was obtained in Fig 6(b) for the N×water interaction, and the maximum SSC/TA ratio was 17.80 at 22.22 g/plant Ca (NO_3_)_2_⋅4H_2_O and 10.58 L/plant water (Fig 6(b)). For the P×water interaction, the SSC/TA ratio slowly increased and then rapidly decreased with increasing P fertilizer and water, and the maximum SSC/TA ratio was 17.64 at 1.67 g/plant NaH_2_PO_4_ and 10.58 L/plant water (Fig 6(c)).

**Fig 6.**
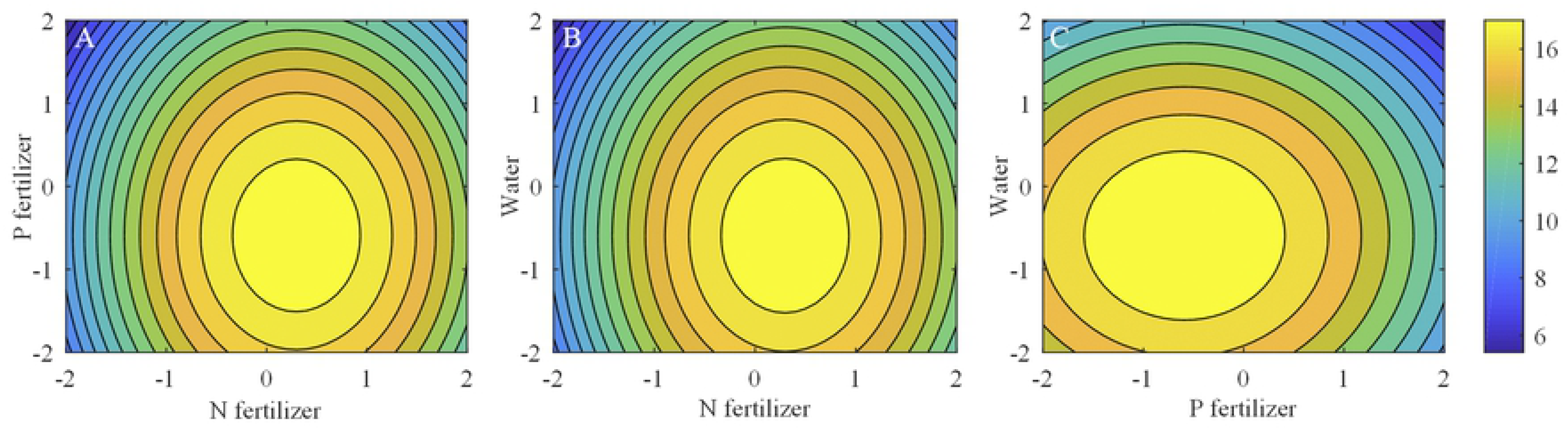
Effects of the interaction among N, P, and K fertilizers and water on the SSC/TA ratio. (A) N-P interaction effect on SSC/TA ratio with water at 0 level. (B) N-water interaction effect on SSC/TA ratio with P fertilizer at 0 level. (C) P-water interaction effect on SSC/TA ratio with P fertilizer at 0 level.

Frequency analysis was performed to obtain the optimal fertilization combination for preferable SSC/TA ratio (Table 8). Among 625 kinds of fertilization combinations, 47 combinations of the three factors were available with strawberry fruit’s SSC/TA ratio varying between 8.5 and 14. The 99% confidence interval for N, P, and water levels were 0.101–0.962, −0.663– 0.407, and −0.650–0.437, respectively. Thus, when applying 21.38–24.99 g/plant Ca (NO_3_)_2_⋅4H_2_O, 1.64–2.04 g/plant NaH_2_PO_4_, and 10.44–13.05 L/plant water, the fruit SSC/TA ratio will reach 8.5–14 with a probability of 99%.

**Table 8.** Frequency analysis of N, P, and water for strawberry fruit’s SSC/TA ratio of 8.5–14.

| Levels | N fertilizer |  | P fertilizer |  | Water |  |
| --- | --- | --- | --- | --- | --- | --- |
|  | Sets Times | Frequency | Sets Times | Frequency | Sets Times | Frequency |
| –2 | 0 | 0.00 | 12 | 0.26 | 12 | 0.26 |
| –1 | 12 | 0.26 | 8 | 0.17 | 8 | 0.17 |
| 0 | 11 | 0.23 | 8 | 0.17 | 8 | 0.17 |
| 1 | 11 | 0.23 | 12 | 0.26 | 11 | 0.23 |
| 2 | 13 | 0.28 | 7 | 0.15 | 8 | 0.17 |
| Total | 47 | 1 | 47 | 1 | 47 | 1 |
| Average | 0.532 |  | –0.128 |  | –0.106 |  |
| Standard error | 0.167 |  | 0.208 |  | 0.211 |  |
| 99% confidence interval | 0.101–0.962 |  | –0.663–0.407 |  | –0.650–0.437 |  |
| Optimal Fertilization (g/plant) | 21.38–24.99 |  | 1.64–2.04 |  | 10.44–13.05 |  |

### Optimal fertilization combination for both high yield and best quality

In accordance with the intersection calculation of the optimal fertilization combination for high yield and best quality, the best fertilization combinations for high yield (more than 110 g/plant) and best fruit SSC/TA ratio (8.5–14) were 22.28–24.61 g/plant Ca (NO_3_)_2_⋅4H_2_O, 1.75–2.03 g/plant NaH_2_PO_4_, 12.41–13.91 g/plant K_2_SO_4_, and 12.00–13.05 L/plant water.

## Discussion

Currently, market intermediaries pay considerable attention to fruit quality to enhance profits by meeting consumers’ preferences of sweetness [11,42], and the SSC/TA ratio has been widely used as a reliable predictor to evaluate the strawberry flavor of sweetness and sourness. Strawberries are sweeter if their SSC/TA ratio is high than if their SSC/TA ratio is low [12,15,28,43,44]. The minimum SSC/TA ratio for acceptable flavor is 8.75 [45], and people prefer the taste of cultivars ‘Clery’ and ‘Daroyal’, which have high SSC/TA ratios of 9.66 and 9.26, respectively [46]. The cultivar ‘NCS 10-156’ has an SSC/TA ratio of 11.6, and it is believed to be more suitable for sale than other cultivars [47]; all these SSC/TA ratios are consistent with the typical range (8.5–14) for strawberries with optimal fruit quality [48,49]. In general, the SSC/TA ratio is an important parameter to evaluate fruit quality for strawberry production [50,51]. Therefore, the SSC/TA ratio was the focus of this study.

Previous studies have shown that N, P, K, and water have significant effects on the yield and fruit quality of strawberry [27,52–55]. A quadratic regression orthogonal rotation combination experimental design was used to investigate the optimal fertilization and water combination for high strawberry yield and best fruit quality with optimal SSC/TA ratio. In the present work, N, P, and water had a significant effect on yield and SSC/TA ratio, whereas K had a significant on yield only. Except for the interaction between N and K having a significant effect on yield, the other interactions among the four factors had no significant effect on yield and SSC/TA ratio. The effects of the four factors on yield and SSC/TA ratio were ranked as N>water>K>P and N>P>water>K, respectively. N was the most important factor among the four factors that had a significant effect on yield and SSC/TA ratio. Thus, N was the key factor in determining the fruit yield and quality. By contrast, when application levels were above 0, P and water had a significant negative effect on the SSC/TA ratio; this result was consistent with findings of previous studies [27,56]. Excessive P suppresses SSC production and promotes TA formation, and excessive water reduces the fruits’ sweetness perception.

The combined application of fertilizer and water should be optimized based on the interaction analysis in the present study. The yield and SSC/TA ratio increased and then decreased as two factors increased but the two other factors were fixed at 0. These trends indicated maximum or optimal values of yield and SSC/TA ratio.

The optimal fertilizer and water combination for high yield (>110 g/plant) and best fruit quality (SSC/TA ratio of 8.5–14) was achieved by using a quadratic regression orthogonal rotation combination experimental design and variance analysis. The optimal fertilizer and water combination was found to be 22.28–24.61 g/plant Ca (NO_3_)_2_⋅4H_2_O, 1.75–2.03 g/plant NaH_2_PO_4_, 12.41–13.91 g/plant K_2_SO_4_, and 12.00–13.05 L/plant water.

## Conclusion

N was the most important factor on yield and SSC/TA, followed by water and P. N, P, and water significantly influenced yield and SSC/TA, whereas K had a significant effect only on yield. The N×K interaction had a significant effect on yield. However, the other interactions among the four factors showed no significant effects on the yield and SSC/TA. The effects of the four factors on the yield and SSC/TA ratio were ranked as N>water>K>P and N>P>water>K, respectively. The yield and SSC/TA ratio increased and then decreased when NPK fertilizers and water increased. The optimal fertilizer and water combination for high yield (>110 g/plant) and best fruit quality (SSC/TA ratio of 8.5–14) was 22.28–24.61 g/plant Ca (NO_3_)_2_⋅4H_2_O, 1.75–2.03 g/plant NaH_2_PO_4_, 12.41–13.91 g/plant K_2_SO_4_, and 12.00–13.05 L/plant water. The results obtained in this study are believed to be useful for further research on fertilization and water application in crops.

## Acknowledgement

We thank Li Shuaishuai, Wang Hongkang and Mu Yonghang (China Agricultural University, Beijing, China) for their help in the investigation discussion, we are grateful for their helpful suggestions.

## References

1. Zhang J, Balkovič J, Azevedo LB, Skalský R, Bouwman AF, Xu G, et al. Analyzing and modelling the effect of long-term fertilizer management on crop yield and soil organic carbon in China. Sci Total Environ. 2018;627: 361–372. doi:10.1016/j.scitotenv.2018.01.090

2. Zhang X, Xu M, Sun N, Xiong W, Huang S, Wu L. Modelling and predicting crop yield, soil carbon and nitrogen stocks under climate change scenarios with fertiliser management in the North China Plain. Geoderma. 2016;265: 176–186. doi:10.1016/j.geoderma.2015.11.027

3. Sauer T, Havlík P, Schneider UA, Schmid E, Kindermann G, Obersteiner M. Agriculture and resource availability in a changing world: The role of irrigation. Water Resour Res. 2010. doi:10.1029/2009WR007729

4. Lu Y, Song S, Wang R, Liu Z, Meng J, Sweetman AJ, et al. Impacts of soil and water pollution on food safety and health risks in China. Environ Int. 2015;77: 5–15. doi:10.1016/j.envint.2014.12.010

5. Zhou K, Sui Y, Xu X, Zhang J, Chen Y, Hou M, et al. The effects of biochar addition on phosphorus transfer and water utilization efficiency in a vegetable field in Northeast China. Agric Water Manag. 2018;210: 324–329. doi:10.1016/j.agwat.2018.08.007

6. Pan J, Liu Y, Zhong X, Lampayan RM, Singleton GR, Huang N, et al. Grain yield, water productivity and nitrogen use efficiency of rice under different water management and fertilizer-N inputs in South China. Agric Water Manag. 2017;184: 191–200. doi:10.1016/j.agwat.2017.01.013

7. FAO. FAOSTAT-DATA-CROPS. [cited 26 Nov 2018]. Available: http://www.fao.org/faostat/en/#data/QC

8. Lado J, Vicente E, Manzzioni A, Aresb G. Application of a check-all-that-apply question for the evaluation of strawberry cultivars from a breeding program. J Sci Food Agric. 2010;90: 2268–2275. doi:10.1002/jsfa.4081

9. Ulrich D, Olbricht K. Searching for chemical and sensory parameters for flavor enhancement in strawberry breeding. Acta Hortic. 2017;1156: 653–659. doi:10.17660/ActaHortic.2017.1156.95

10. Keutgen A, Pawelzik E. Modifications of taste-relevant compounds in strawberry fruit under NaCl salinity. Food Chem. 2007;105: 1487–1494. doi:10.1016/j.foodchem.2007.05.033

11. Cao F, Guan C, Dai H, Li X, Zhang Z. Soluble solids content is positively correlated with phosphorus content in ripening strawberry fruits. Sci Hortic (Amsterdam). 2015;195: 183–187. doi:10.1016/j.scienta.2015.09.018

12. Ulrich D, Olbricht K. A search for the ideal flavor of strawberry - Comparison of consumer acceptance and metabolite patterns in Fragaria x ananassa Duch. J Appl Bot Food Qual. 2016;89: 223-+. doi:10.5073/Jabfq.2016.089.029

13. Crisosto CH, Garner D, Crisosto GM, Bowerman E. Increasing “Blackamber” plum (Prunus salicina Lindell) consumer acceptance. Postharvest Biol Technol. 2004;34: 237–244. doi:10.1016/j.postharvbio.2004.06.003

14. Alavoine F, Crochon M. Taste quality of strawberry. Acta Hortic. 1989;265: 449–452. doi:10.17660/ActaHortic.1989.265.68

15. Gunness P, Kravchuk O, Nottingham SM, D’Arcy BR, Gidley MJ. Sensory analysis of individual strawberry fruit and comparison with instrumental analysis. Postharvest Biol Technol. 2009;52: 164–172. doi:10.1016/j.postharvbio.2008.11.006

16. Li H, Lascano RJ. Deficit irrigation for enhancing sustainable water use: Comparison of cotton nitrogen uptake and prediction of lint yield in a multivariate autoregressive state-space model. Environ Exp Bot. 2011;71: 224–231. doi:10.1016/j.envexpbot.2010.12.007

17. Dordas CA, Sioulas C. Safflower yield, chlorophyll content, photosynthesis, and water use efficiency response to nitrogen fertilization under rainfed conditions. Ind Crops Prod. 2008;27: 75–85. doi:10.1016/j.indcrop.2007.07.020

18. Deng X, Woodward FI. The growth and yield responses of Fragaria ananassa to elevated CO2 and N supply. Ann Bot. 1998;81: 67–71. doi:10.1006/anbo.1997.0535

19. Tsialtas JT, Maslaris N. Effect of N fertilization rate on sugar yield and non-sugar impurities of sugar beets (Beta vulgaris) grown under Mediterranean conditions. J Agron Crop Sci. 2005;191: 330–339. doi:10.1111/j.1439-037X.2005.00161.x

20. Li H, Li T, Fu G, Hu K. How strawberry plants cope with limited phosphorus supply: Nursery-crop formation and phosphorus and nitrogen uptake dynamics. J Plant Nutr Soil Sci. 2014;177: 260–270. doi:10.1002/jpln.201200654

21. Grant CA, N FD, Tomasiewiz DJ, Jonston AM. The importance of early season phosphorus nutrition. Can J Plant Sci. 2001;81: 60–73. doi:10.4141/P00-093

22. Oosterhuis DM, Loka DA, Kawakami EM, Pettigrew WT. The physiology of potassium in crop production. Advances in Agronomy. Elsevier; 2014. doi:10.1016/B978-0-12-800132-5.00003-1

23. Pettigrew WT. Potassium influences on yield and quality production for maize, wheat, soybean and cotton. Physiol Plant. 2008;133: 670–681. doi:10.1111/j.1399-3054.2008.01073.x

24. Tsialtas IT, Shabala S, Baxevanos D, Matsi T. Effect of potassium fertilization on leaf physiology, fiber yield and quality in cotton (Gossypium hirsutum L.) under irrigated Mediterranean conditions. F Crop Res. 2016;193: 94–103. doi:10.1016/j.fcr.2016.03.010

25. Grant OM, Johnson AW, Davies MJ, James CM, Simpson DW. Physiological and morphological diversity of cultivated strawberry (Fragaria × ananassa) in response to water deficit. Environ Exp Bot. 2010;68: 264–272. doi:10.1016/j.envexpbot.2010.01.008

26. Martínez-Ferri E, Soria C, Ariza MT, Medina JJ, Miranda L, Domíguez P, et al. Water relations, growth and physiological response of seven strawberry cultivars (Fragaria×ananassa Duch.) to different water availability. Agric Water Manag. 2016;164: 73–82. doi:10.1016/j.agwat.2015.08.014

27. Weber N, Zupanc V, Jakopic J, Veberic R, Mikulic-Petkovsek M, Stampar F. Influence of deficit irrigation on strawberry (Fragaria × ananassa Duch.) fruit quality. J Sci Food Agric. 2017;97: 849–857. doi:10.1002/jsfa.7806

28. Akhtar I, Rab A. Effect of Irrigation Intervals on the Quality and Storage Performance of Strawberry Fruit. J Anim Plant Sci. 2015;25: 669–678.

29. Ener SŞ. Effects of Genotype and Fertilization on Fruit Quality in Several Harvesting Periods of Organic Strawberry Plantation. 2016;5: 252–256.

30. McArthur DAJ, Eaton GW. Strawberry yield response to fertilizer, paclobutrazol and chlormequat. Sci Hortic (Amsterdam). 1988;34: 33–45. doi:10.1016/0304-4238(88)90073-8

31. Pokhrel B, Laursen KH, Petersen KK. Yield, Quality, and Nutrient Concentrations of Strawberry (Fragaria × ananassa Duch. cv. Sonata) Grown with Different Organic Fertilizer Strategies. J Agric Food Chem. 2015;63: 5578–5586. doi:10.1021/acs.jafc.5b01366

32. Sønsteby A, Opstad N, Myrheim U, Heide OM. Interaction of short day and timing of nitrogen fertilization on growth and flowering of “Korona” strawberry (Fragaria × ananassa Duch.). Sci Hortic (Amsterdam). 2009;123: 204–209. doi:10.1016/j.scienta.2009.08.009

33. Wang B, Lai T, Huang QW, Yang XM, Shen QR. Effect of N Fertilizers on Root Growth and Endogenous Hormones in Strawberry Project supported by the National High Technology Research and Development Program (863 Program) of China (No. 2004AA246080) and the Program for the Development of High-Tech Indu. Pedosphere. 2009;19: 86–95. doi:10.1016/S1002-0160(08)60087-9

34. Bottoms TG. Nitrogen Management and Water Quality Protection in California Lettuce and Strawberry Production by THOMAS GLENN BOTTOMS B. S. (California Polytechnic State University, San Luis Obispo) 2008 DISSERTATION Submitted in partial satisfaction of the requi. University of California, Davis. 2013.

35. Xu D, Du S, van Willigenburg LG. Optimal control of Chinese solar greenhouse cultivation. Biosyst Eng. 2018;171: 205–219. doi:10.1016/j.biosystemseng.2018.05.002

36. Ahamed MS, Guo H, Tanino K. Development of a thermal model for simulation of supplemental heating requirements in Chinese-style solar greenhouses. Comput Electron Agric. 2018;150: 235–244. doi:10.1016/j.compag.2018.04.025

37. Xing M, Sun K, Liu Q, Pan L, Tu K. Development of Novel Electronic Nose Applied for Strawberry Freshness Detection during Storage. Int J Food Eng. 2018;0: 1–15. doi:10.1515/ijfe-2018-0111

38. Zhang C, Guo C, Liu F, Kong W, He Y, Lou B. Hyperspectral imaging analysis for ripeness evaluation of strawberry with support vector machine. J Food Eng. 2016;179: 11–18. doi:10.1016/j.jfoodeng.2016.01.002

39. Shao X, Wang H, Xu F, Cheng S. Effects and possible mechanisms of tea tree oil vapor treatment on the main disease in postharvest strawberry fruit. Postharvest Biol Technol. 2013;77: 94–101. doi:10.1016/j.postharvbio.2012.11.010

40. Liu C, Zheng H, Sheng K, Liu W, Zheng L. Effects of melatonin treatment on the postharvest quality of strawberry fruit. Postharvest Biol Technol. 2018;139: 47–55. doi:10.1016/j.postharvbio.2018.01.016

41. Yuan Y, Tan L, Xu Y, Dong J, Zhao Y, Yuan Y. Optimization of Processing Parameters for lettuce Vacuum Osmotic Dehydration Using Response Surface Methodology. Polish J Food Nutr Sci. 2018;68: 15–23. doi:10.1515/pjfns-2017-0013

42. Gallardo RK, Li H, McCracken V, Yue C, Luby J, McFerson JR. Market Intermediaries’ Willingness to Pay for Apple, Peach, Cherry, and Strawberry Quality Attributes. Agribusiness. 2014;0: 1–22. doi:10.1002/agr.21396

43. Cordenunsi BR, Nascimento JRO, Lajolo FM. Physico-chemical changes related to quality of five strawberry fruit cultivars during cool-storage. Food Chem. 2003;83: 167–173. doi:10.1016/S0308-8146(03)00059-1

44. Weissinger H, Eggbauer R, Steiner I, Spornberger A, Steffek R, Altenburger J, et al. Yield and fruit quality parameters of new early-ripening strawberry cultivars in organic growing on a highly Verticillium -infested site. 14th Int Conf Org fruit Grow. 2010; 243–249.

45. Pelayo-Zaldívar C, Ebeler SE, Kader AA. CULTIVAR AND HARVEST DATE EFFECTS ON FLAVOR AND OTHER QUALITY ATTRIBUTES OF CALIFORNIA STRAWBERRIES. J Food Qual. 2005;28: 78–97.

46. Voca S, Dobricevic N, Dragovic-Uzelac V, Duralija B, Druzic J, Cmelik ZSB. Fruit Quality of New Early Ripening Strawberry Cultivars in Croatia. Food Technol Biotechnol. 2008;46: 292–298.

47. Perkins-Veazie P, Pattison J, Fernandez G, Ma G. Fruit Quality and Composition of Two Advanced North Carolina Strawberry Selections. Int J Fruit Sci. 2016;16: 220–227. doi:10.1080/15538362.2016.1219289

48. Montero TM, Mollá EM, Esteban RM, López-Andréu FJ. Quality attributes of strawberry during ripening. Sci Hortic (Amsterdam). 1996;65: 239–250. doi:10.1016/0304-4238(96)00892-8

49. Kafkas E, Koşar M, Paydaş S, Kafkas S, Başer KHC. Quality characteristics of strawberry genotypes at different maturation stages. Food Chem. 2007;100: 1229–1236. doi:10.1016/j.foodchem.2005.12.005

50. Rutkowski KP, Kruczynska DE, Zurawicz E. Quality and shelf life of strawberry cultivars in Poland. V Int Strawb Symp 708. 2004; 329–332.

51. Wang Q, Tury E, Rekika D, Thérèse Charles M, Tsao R, Hao YJ, et al. Agronomic characteristics and chemical composition of newly developed day-neutral Strawberry lines by agriculture and agri-food Canada. Int J Food Prop. 2010;13: 1234–1243. doi:10.1080/10942910903013415

52. Cronje RB, Sivakumar D, Mostert PG, Korsten L. Effect of different preharvest treatment regimes on fruit quality of litchi cultivar “Maritius.” J Plant Nutr. 2009;32: 19–29. doi:10.1080/01904160802530987

53. Milošević TM, Glišić IP, Glišić IS, Milošević NT. Cane properties, yield, berry quality attributes and leaf nutrient composition of blackberry as affected by different fertilization regimes. Sci Hortic (Amsterdam). 2018;227: 48–56. doi:10.1016/j.scienta.2017.09.013

54. Chelpiński P, Skupień K, Ochmian I. Effect of fertilization on yield and quality of cultivar kent strawberry fruit. J Elem. 2010;15: 251–257.

55. Cardeñosa V, Medrano E, Lorenzo P, Sánchez-Guerrero MC, Cuevas F, Pradas I, et al. Effects of salinity and nitrogen supply on the quality and health-related compounds of strawberry fruits (Fragaria × ananassa cv. Primoris). J Sci Food Agric. 2015;95: 2924–2930. doi:10.1002/jsfa.7034

56. Moor U, Põldma P, Tõnutare T, Karp K, Starast M, Vool E. Effect of phosphite fertilization on growth, yield and fruit composition of strawberries. Sci Hortic (Amsterdam). 2009;119: 264–269. doi:10.1016/j.scienta.2008.08.005

